# OME-NGFF: scalable format strategies for interoperable bioimaging data

**DOI:** 10.1101/2021.03.31.437929

**Authors:** Josh Moore, Chris Allan, Sebastien Besson, Jean-Marie Burel, Erin Diel, David Gault, Kevin Kozlowski, Dominik Lindner, Melissa Linkert, Trevor Manz, Will Moore, Constantin Pape, Christian Tischer, Jason R. Swedlow

## Abstract

Biological imaging is one of the most innovative fields in the modern biological sciences. New imaging modalities, probes, and analysis tools appear every few months and often prove decisive for enabling new directions in scientific discovery. One feature of this dynamic field is the need to capture new types of data and data structures. While there is a strong drive to make scientific data Findable, Accessible, Interoperable and Reproducible (FAIR ^1^), the rapid rate of innovation in imaging impedes the unification and adoption of standardized data formats. Despite this, the opportunities for sharing and integrating bioimaging data and, in particular, linking these data to other “omics” datasets have never been greater. Therefore, to every extent possible, increasing “FAIRness” of bioimaging data is critical for maximizing scientific value, as well as for promoting openness and integrity.

In the absence of a common, FAIR format, two approaches have emerged to provide access to bioimaging data: translation and conversion. On-the-fly translation produces a transient representation of bioimage metadata and binary data but must be repeated on each use. In contrast, conversion produces a permanent copy of the data, ideally in an open format that makes the data more accessible and improves performance and parallelization in reads and writes. Both approaches have been implemented successfully in the bioimaging community but both have limitations. At cloud-scale, those shortcomings limit scientific analysis and the sharing of results. We introduce here next-generation file formats (NGFF) as a solution to these challenges.

## Limits of On-the-fly Translation

One classically successful approach to provide unified access to bioimaging data uses a library that dynamically translates bioimage metadata and binary data from the innumerable proprietary file formats (PFFs) that exist in this domain (Figure 1a). This strategy is provided by several open source solutions, including the authors’ Bio-Formats ^2^, a Java-based library that supports >150 PFFs; OpenSlide ^3^, a C++ library that focuses on PFFs used in whole slide imaging (WSI); and aicsimageio ^4^, a Python library that wraps vendor libraries for simplified numpy access. Translating data in real-time has worked well for many applications, and such translation libraries have become the reference APIs in their fields. However, large, public data resources like the Allen Cell Explorer (https://www.allencell.org/), Image Data Resource ^5^, Systems Science of Biological Dynamics Database ^6^, and others have revealed the fundamental bottlenecks created by the computational cost and time required for repeated translation of massive collections of PFFs. The same problem occurs in data hungry applications like machine learning (ML), where the cost of real-time translation precludes the use of even larger, more richly annotated datasets. Furthermore, the outputs of these applications remain fundamentally isolated from the original data when either the complexity or licensing of PFFs prevents the writing of further analytical metadata in the same format. Finally, individual dataset volumes have grown with the advent and popularization of imaging modalities that support large tissue samples, such as digital pathology, light sheet microscopy (LSM), and focused on beam-scanning electron microscopy. In these applications, efficient, high performance data access requires multi-resolution representations (often referred to as pyramidal data) that enable zoomable visualization and selectable levels of resolution for interactive navigation and scalable analysis. Providing multi-resolution support across >150 PFFs is simply not practical nor computationally effective. In short, for the applications that are becoming strategic opportunities for new directions in bioimaging, real-time translation no longer scales.

**Figure 1.**
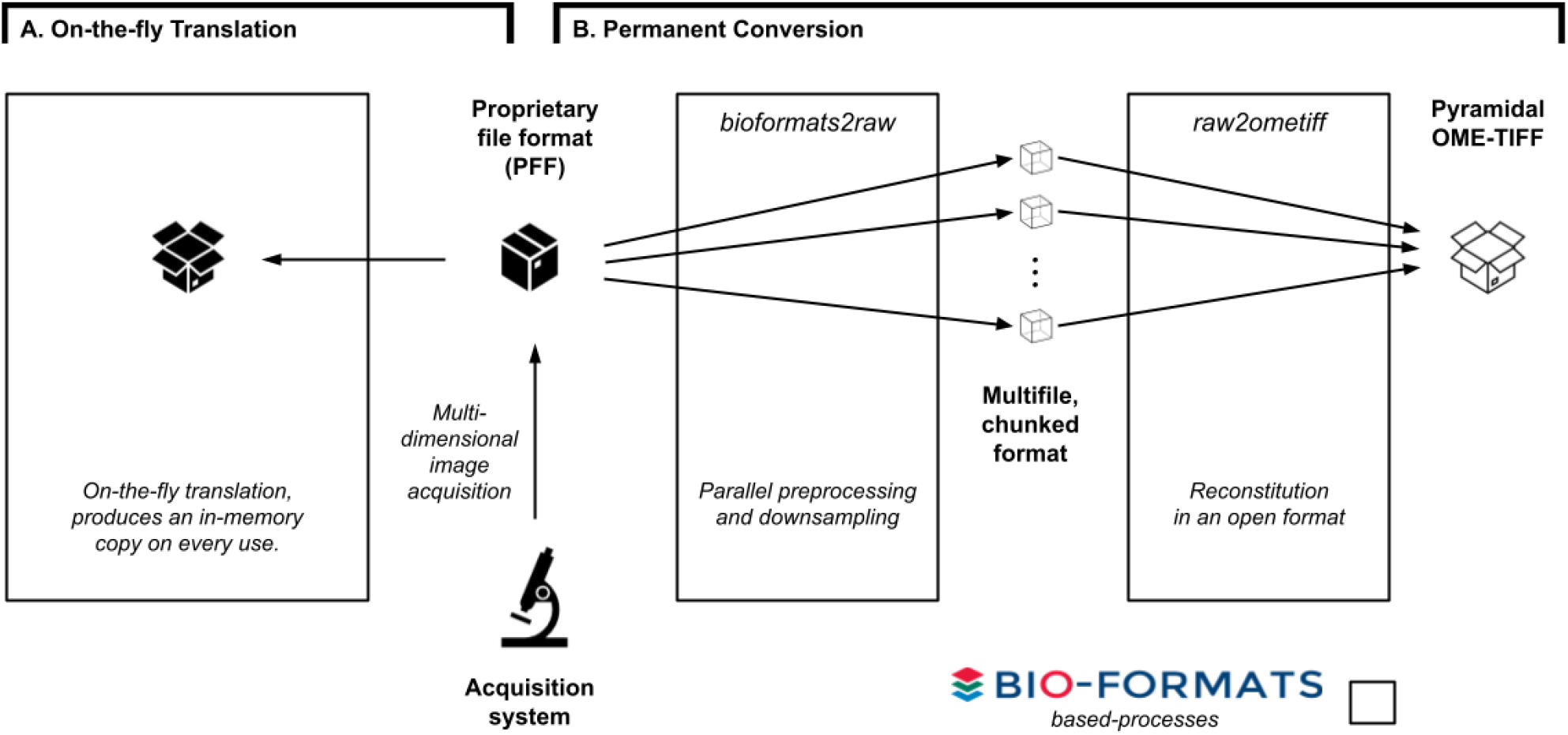
Conversion tools provide an alternative to continual, on-the-fly translation of PFFs. *Figure shows workflows for file format conversion. A. The classical approach to access images produced by an acquisition system is to use a library like Bio-Formats to translate the proprietary file format (PFF) and produce an in-memory copy of the imaging data on-the-fly. This translation needs to be repeated on every use. B. With the existence of open, community-supported formats, converting PFFs becomes the most cost-efficient method for long-term storage and sharing of microscopy data.* bioformats2raw *and* raw2ometiff *parallelize the creation of an open format, OME-TIFF, by using an intermediate format consisting of many, individual files each with one chunk of the original image data.*

## Permanent Conversion

A distinct approach -- converting data from PFFs to a common, well-defined format -- solves the computational demands of repeated and real-time image translation but requires a format that has broad application and utility, long-term stability and multiple open-source implementations and the support of the community. Some success has been achieved with OME-TIFF, a 2D multi-resolution image format that captures acquisition metadata as OME-XML in the TIFF header ^2,7,8^. Reference software implementations are available in Java (https://github.com/ome/bioformats/), C++ (https://gitlab.com/codelibre/ome/ome-files-cpp) and Python (e.g., https://github.com/AllenCellModeling/aicsimageio, https://github.com/apeer-micro/apeer-ometiff-library, https://github.com/cgohlke/tifffile). OME-TIFF is supported by several commercial imaging companies (see https://www.openmicroscopy.org/commercial-partners/) and is the recommended format for public data projects like Image Data Resource (IDR) or Allen Institute of Cell Science, making their data available from https://open.quiltdata.com/b/allencell/.

As our and others’ use of existing tools for conversion to OME-TIFF grew, TIFF’s linear binary layout became a bottleneck. Larger files took increasingly long to write. This problem was most obvious in projects that required the conversion of large numbers of whole slide images from PFFs to OME-TIFF for use in data lakes that are used for AI training sets (https://pathlake.org; https://icaird.com). The need for a scalable conversion motivated our development of two tools, *bioformats2raw (https://github.com/glencoesoftware/bioformats2raw)* and *raw2ometiff (https://github.com/glencoesoftware/raw2ometiff)*. Together they provide a parallel pipeline using Bio-Formats to convert any supported PFF into multi-resolution OME-TIFF. This is achieved by breaking images into atomic “chunks”, writing them independently to disk, and generating subresolutions from them when none are available, whereupon a second process can efficiently write these chunks into TIFF (Figure 1b). With this conversion pipeline, OME-TIFF becomes a performant solution for domains handling larger planar data, for example whole slide images in digital pathology.

## Multi-dimensional Images

A fundamental issue with OME-TIFF is that though it supports 5D images, its binary data access is limited to 2D tiles. While storage of individual planes of image data that encompass hundreds of GBytes and dozens of pyramidal resolutions of data is possible, performance suffers when scaling to multi-dimensional, multi-TByte datasets, e.g., LSM datasets. In fact, in the benchmark below the LSM-like test data formatted in TIFF consumed too much memory for completing the benchmark.

A possible solution to the dimensionality issue is HDF5, a multi-dimensional data format that internally supports chunk-based access ^9^. Several open HDF5 bioimage file formats have been designed and implemented ^10–13^ and libraries exist for these various formats in several of the major programming languages. The HDF5-based BigDataViewer file format ^11^ has proven to be quite powerful for the LSM community, as it provides a convenient integrated format and the chunking required for interactive visualization of the large 3D timelapse datasets produced by LSM. Oxford Instruments have released an alternative open HDF5-based implementation that is widely used and includes a format specification (https://imaris.oxinst.com/support/imaris-file-format) as well as an open reference implementation, ImarisWriter ^14^. These formats have performed well for interactive visualization and analysis, but there remain limitations for processing and integration of large numbers of HDF5 datasets. Parallel writing of HDF5 files is only fully supported in high performance, specialised parallel computing environments where technologies such as MPI (“message passing interface”) are available. Also, unlike TIFF, HDF5 files do not inherently have a specification for image data, so each HDF5 implementation ends up being another file format, growing the number of PFFs.

## Cloud Performance

An issue with both TIFF and HDF5 is that any format must eventually be stored on a hard-drive or other permanent memory structure. Filesystems have been the workhorse of the imaging community since its inception. They enable low-latency, “random access” to large binary data files. This speed is an underlying assumption of most visualization and analysis applications, but filesystems are relatively expensive and their complexity comes with relatively high maintenance costs. With the ascendance of cloud computing, an alternative is to use a loosely-defined storage technology (often called “object storage”) that treats individual files as distinct immutable objects. Object storage provides relatively simple read and write procedures that transfer whole objects (often called “chunks”). Each object is stored redundantly across multiple servers, which offers improved parallelization and scalability in exchange for increased access latency. To make use of the advantages of object storage, a modern format, accommodating contemporary dataset volumes and dimensionality, is needed that does not require the whole binary structure to be accessed as a single monolithic block on disk or in the cloud. While a monolithic strategy works for smaller files, it is fundamentally limiting for multi-TByte, multi-dimensional and multi-modal imaging datasets. These datasets are so large that they cannot be completely transferred between computers, locations or public resources, even if sufficient storage were available, which is rarely the case. A new form of data access is needed in bioimaging.

While the TIFF and HDF5 implementations remain valuable in their respective domains, their inability to cover all use cases is illustrative of the pitfalls of format standardization. Creating a data format standard requires equal consideration for performance, usability, and structure, with a balance of community-driven specification and extensibility. Historically, OME-TIFF and OME-XML were highly specified but lacked optimal adaptability to novel data volumes and high dimensionality, while HDF5 was highly extensible and thereby suffered a branching into multiple PFFs. Improved performance motivated early adoption, especially given high usability, but providing a clear structure for binary data and metadata is essential to yield a cohesive landscape of new tools, rather than a divergence of format variants. Extensibility, however, is necessary for adoption by new domains or vendors and integration with novel analytical approaches that were not considered at the point of initial specification.

## Next-generation file formats

To address these challenges, we have begun preparing a new generation of binary file formats for bioimaging. The work is based on the tenet that, given the diversity of the field of bioimaging, no single data structure or file format will address all use cases, programming environments and technology domains. For instance the requirements of an imaging technology manufacturer, which needs a file format with optimised write performance, are very different from those of a computational scientist building new ML technologies or a public data repository that has to publish and serve many millions to billions of image datasets.

As we investigated candidates for the intermediate multifile, “chunked” representation for parallelizing the *bioformats2raw* and *raw2ometiff* conversion pipeline (see Figure 1), we realized that these data structures could complement existing TIFF- and HDF5-based workflows and enable new types of parallel bioimaging use cases. We collectively refer to these formats as “next generation file formats” (NGFFs). We have focused on two very similar, open source strategies for laying out binary data. N5 (https://github.com/saalfeldlab/n5) is a binary data format that uses embedded directories and defined multi-dimensional chunking to provide fast, cloud-competent image data storage. N5 was developed out of the Fiji community ^15^ and there are now several examples of public datasets in N5. Zarr (https://github.com/zarr-developers) follows a nearly identical strategy of storing chunks in individual files across directories and was originally adopted for handling genomic and geospatial data, similarly with a number of datasets available publicly ^16^. Since 2018, the two communities have worked together (https://github.com/zarr-developers/community/issues/1#issue-463966208) to unify the two formats to maximize re-use. Table 1 summarizes the overall added value of the NGFFs.

**Table 1.** Characteristics of three binary containers for imaging data. To accomplish this idea, we have defined an image specification for multiscale NGFFs on top of N5 and Zarr. We have also built implementations of the specification, demonstrating the usability and performance of these formats. *bioformats2raw* can be used for writing NGFF formats from standalone Java applications and *omero-cli-zarr* is available for exporting from OMERO ^17^. Reading is implemented in *ome-zarr-py* which has been integrated into the *napari* viewer ^18^, in *Fiji* via the *MoBIE* plugin ^19^, and finally via Viv-based *vizarr* for access in the browser ^20^. Each additional implementation enables a different set of users to remotely access data without the overhead of PFF translation. Permissively-licensed datasets from IDR have been converted into Zarr and stored in an S3 object storage bucket for public consumption (Figure 2).

|  | Maturity | Dimensionality | Scalability |
| --- | --- | --- | --- |
| TIFF | Ubiquitous | 2D tiles | Poor |
| HDF5 | Well-supported | 3D+ chunks | Limited parallel access |
| Zarr/N5 | Next-generation | 3D+ chunks | Fully parallel |

**Figure 2.**
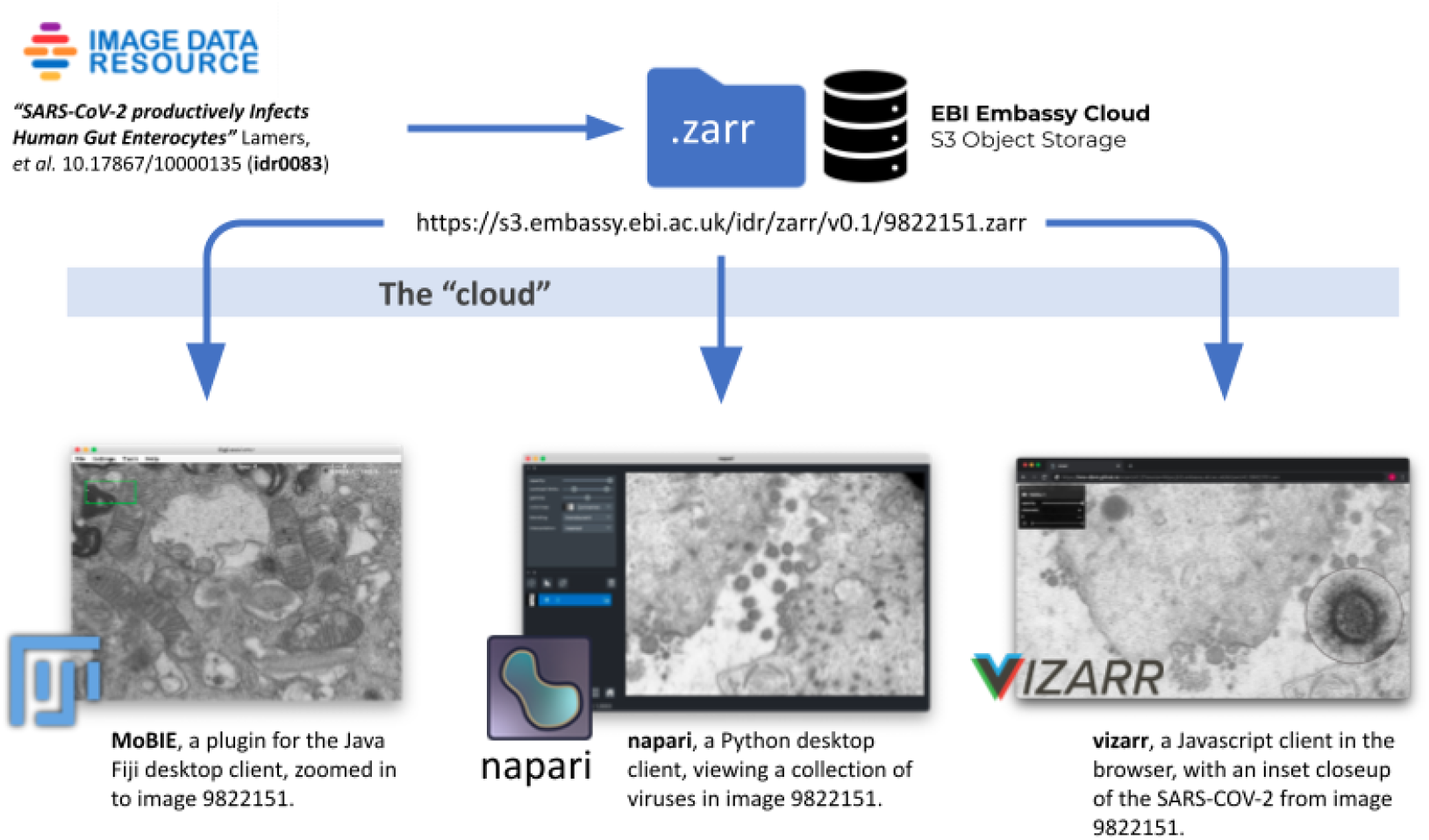
Maximizing re-use by allowing popular tools to access bioimaging data in the cloud. *An example of using NGFFs for promoting the distribution of public image datasets. Selection of current tools streaming different portions of the same SARS-COV-2 virus image at various resolutions directly from S3 storage at the European Bioinformatics Institute (EBI). Original data from Lamers et al. DOI:10.1126/science.abc1669 available in IDR at DOI:10.17867/10000135*.

It should be noted that although this set of formats provides an initial basis for development, as the needs of the community are catalogued, more sophisticated formats like TileDB Embedded may join this list of supported candidates. TileDB Embedded uses a database-like storage engine based on sparse and dense multi-dimensional arrays to service chunks of data and as it is an open source C++ run-time library, data can be accessed from Java, Python and several other programming languages. However, a long-term commitment to format support by the users, developers, and funders should be demonstrated to avoid a new translation bottleneck.

## Measuring access times

To illustrate the impact of these new binary formats in modern computing environments, we benchmarked the latency of random chunk access across uncompressed datasets representative of two established imaging modalities: large multi-channel two-dimensional images like the ones produced by cyclic immunofluorescence (CycIF) ^21^ and timelapse isotropic volumes typically generated by LSM ^22^. Synthetic HDF5, TIFF and Zarr datasets were generated by first invoking the ImarisWriter, then converting the HDF5-based Imaris files into Zarr with *bioformats2raw*, and finally converting the Zarr to TIFF with *raw2ometiff*. All three datasets along with a 1-byte dummy file for measuring overhead were placed in three types of storage: local disk, a remote server, and in object storage. We measured the reading time of individual chunks for all four file types across the three storage systems, making use of fspec for all access (Figure 3). All code is available for generating the data and running the benchmark in docker or on the AWS cloud.

**Figure 3.**
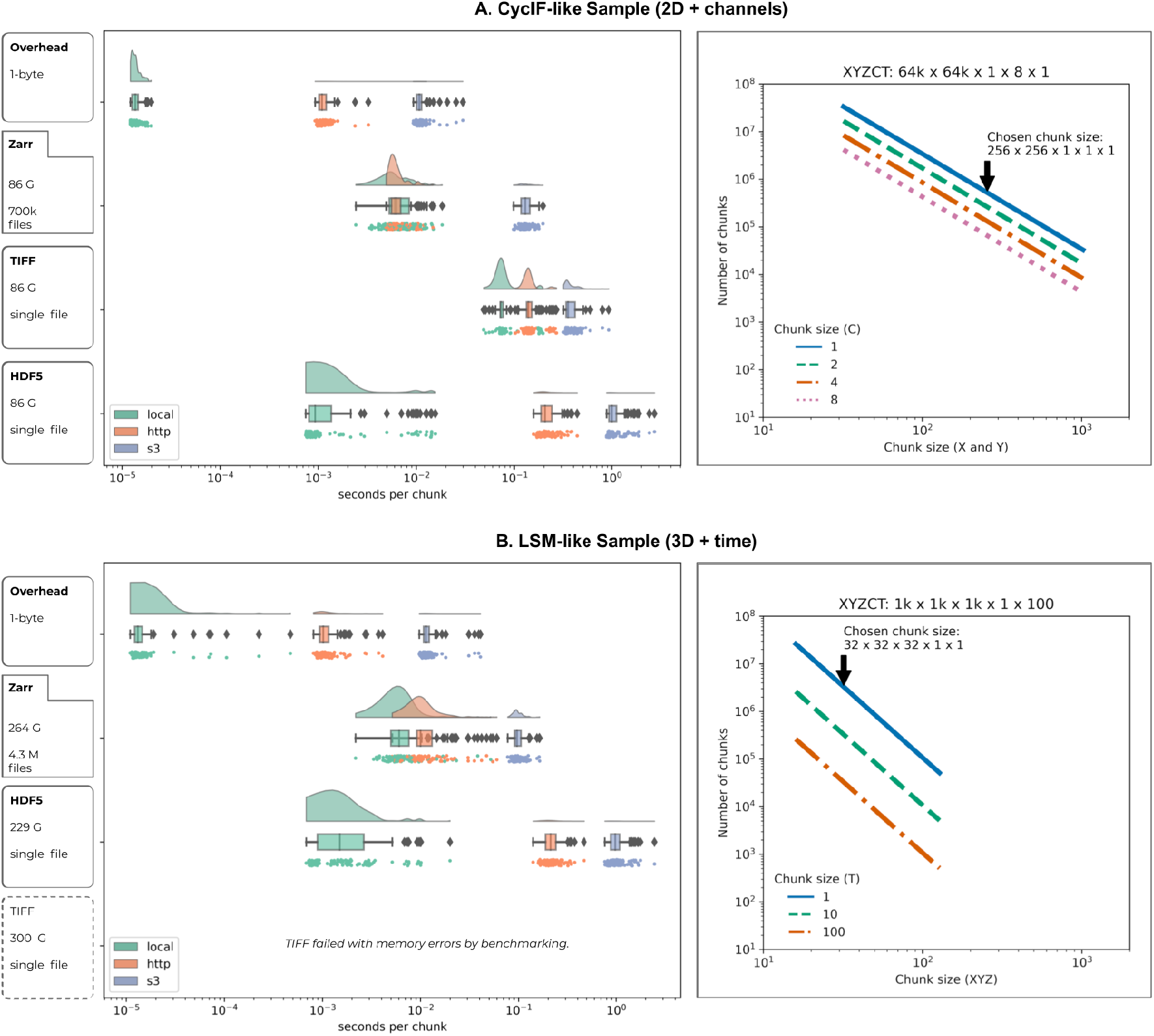
Chunk retrieval time is less sensitive to data location with next-generation file formats. *Random sampling of 100 chunks from synthetically generated, 5D images measures access times for three different formats on the same file system (“local”, green), over HTTP using the nginx web server (“http”, orange),and using Amazon’s proprietary S3 object storage protocol (“s3”, blue) under two scenarios: (A) a whole-slide CycIF imaging dataset with many large planes of data (x=64k, y=64k, c=8) and chunks of 256×256 pixels (128 KB) and (B) a time-lapse LSM dataset with isotropic dimensions (x=1024, y=1024, z=1024, t=100) and chunks of 32×32×32 pixels (64 KB)*. *The rain cloud plots on the left show that as the latency of access grows, access times for monolithic formats like TIFF and HDF5 increase because libraries must seek the appropriate data chunk, whereas NGFF formats like Zarr provide direct access to individual chunks. In the 3D case, the TIFF data was too large to fit into local memory and the benchmark errored*. *The line plots on the right show how the number of chunks needed for an image of the specified dimensions varies with size of the chunk in X and Y (and Z for 3D). Chunk sizes were chosen here to balance chunk size with number of files. All data was stored uncompressed to keep chunk sizes consistent for the random generated data*. *All code for reproducing the runs both locally with docker or Amazon EC2 instances are available in https://github.com/ome/ngff-latency-benchmark*.

We observed that access speeds for NGFF files are similar to HDF5 when the files reside locally (i.e., on the same computer) and both substantially outperform TIFF. We note Kang showed that for most serial access patterns, HDF5 can be significantly faster than Zarr ^23^. Together these results partially explain HDF5’s popularity for desktop analysis and visualization of LSM datasets. But access speeds for NGFF files are at least an order of magnitude faster than HDF5 on object storage. This leads to throughput comparable to local disk access when using parallel reads ^24^, supporting streaming of image data files from remote http-based or cloud-based servers. This speed improvement is achieved by enabling parallelised, chunk-based data access where data is stored in the cloud. This obviates the need for wholesale data download and is especially important for providing performant access to multi-TByte datasets.

A key parameter in overall access times is the size of individual chunks. As chunk sizes decrease, the number of individual chunk files increases rapidly. In this benchmark, we have chosen a compromise between chunk size and number of individual files. This illustrates the primary downside of NGFF formats: as the number of files increases, so does the on-disk complexity with the result that copying the entire fileset between locations now becomes limiting. This is a direct result of optimizing NGFFs for incremental rather than bulk access.

## Interoperable through metadata specifications

Though we have demonstrated the performance benefits of a low-latency streaming format when accessing large multi-dimensional volumes, it is also clear that multiple open bioimaging formats have a role to play in the many use-cases of FAIR data access. Supporting multiple community formats, however, runs the risk of again permanently requiring a translation library. The solution is to guarantee interoperability between the formats themselves. All data in one format should be losslessly convertible to the other, supported formats. The key to achieving this is a re-evaluation of the metadata format that OME has been maintaining for 15 years.

While TIFF provides a standardized specification for defining a 2D-binary image, the extension to accommodate 5D images was only provided via the addition of OME-XML stored in TIFF’s header structure ^2^. The definition of such “alignment” metadata, clearly specifying the relationships between dimensions and measurements, is critical for the interpretation of evolving imaging data. In this modernization of the original XML work, we have added metadata conventions into the metadata structures of the Zarr format to handle multi-dimensional bioimaging data, including high-content screening datasets . We have provided a specification for paired, derived labelled images such as segmentation or classification masks. We believe that the remote streaming capabilities of NGFFs will complement high-throughput ML training and testing workflows, whose outputs can now remain in a common data structure with the original pixel data and metadata.

Metadata for experimental procedures, image acquisition parameters and analytics are another critical component of a bioimaging file format and provide the basis for interoperable data access. The ability to extend and adapt a standard is of particular importance for converting files permanently away from PFFs. Technology developers and commercial manufacturers need clear guidelines and support for incorporating their metadata into and adapting to a community-standard. We have published the current work as a formal specification, following patterns developed by the World Wide Web Consortium that have been successful in other domains, and provided means for public commentary and contribution (https://ngff.openmicroscopy.org/latest/).

A general problem in metadata specifications, however, is that they necessarily focus on a specific domain. While valuable, this can make extension to other domains difficult. An example is the OME-XML specification ^7^ which is focused on fluorescence microscopy, high content screening and digital pathology, and does not provide an easy way for third parties to add support for other modalities or application domains. The redefinition of the OME metadata for NGFFs provides an opportunity to specify conventions that will apply to and support the breadth of formats needed by the community (Figure 4). In collaboration with RIKEN, initial work on a Resource Description Framework (RDF)-based representation of OME metadata has been completed that needs integrating into the NGFF structure ^25–27^.

**Figure 4.**
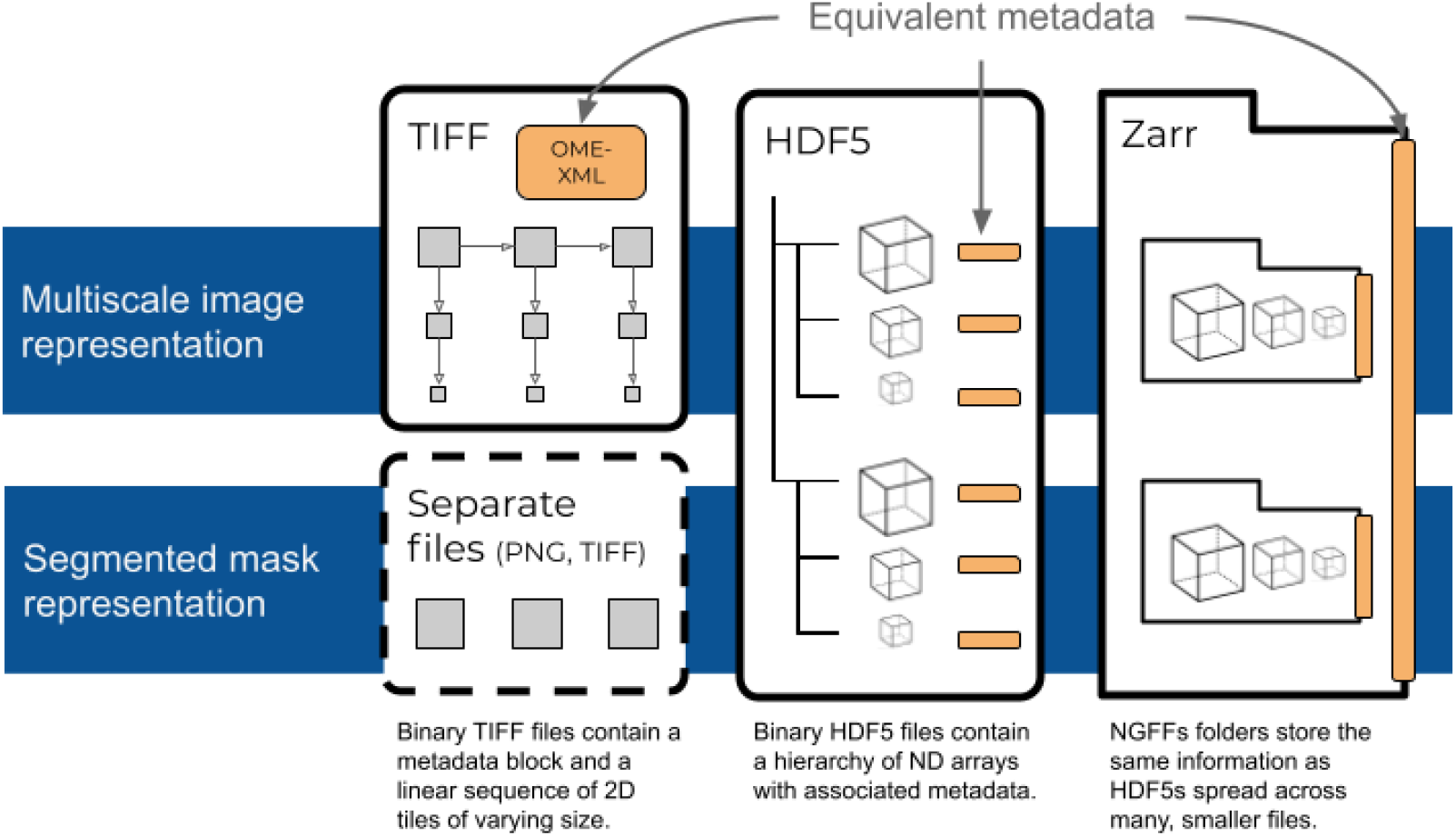
Unification of metadata specifications will allow interoperability between TIFF, HDF5, and Zarr. *Each format is suited to particular use cases. TIFF is ideal for interoperability in digital pathology and other 2-dimensional domains since the format is widely accessible by established open source and proprietary software. In higher-dimensional domains, HDF5 and Zarr are better suited. HDF5 will likely be preferred for local access. If data is intended for sharing in the cloud, Zarr will likely be preferred. High-throughput image analysis will benefit from the lower-latency access to data in HDF5 and Zarr. If original image data is paired with derived representations like pixel or object classification, a shared structure in HDF5 or Zarr is likely the best choice.*

While the rapid pace of innovation in bioimaging continues, the core data structures-- binary pixel data and experimental, acquisition and analytic metadata-- are likely common to most imaging experiments and datasets. Indeed, recent work on minimal bioimaging metadata standards have focused on a collection of core concepts including those from OME that can be extended to support domain- and application-specific requirements. As a participant in the “Recommended Metadata for Biological Images” (Sarkans et al, Nature Methods, in press) effort, we aim to provide open reference software and formats that implement the proposed minimal metadata specification and can be widely adopted and extended by the bioimaging community.

## Outlook: Suite of standard formats

Is it practical to expect that these next generation formats may one day become community-agreed and widely adopted standards? Reducing the explosion of PFFs is essential for enabling data repositories and maintaining the scale at which our community is working, but to date, the community has not been able to agree on any single file format. Any format will be challenged by the level of innovation that has prevented easy adoption of any of the previous candidates.

We assert that this suite of interoperable formats -- TIFF, HDF5, and now low-latency, cloud-capable NGFFs -- will provide the balanced set of trade-offs necessary for the community to converge on, slowing the development of ever more file formats. When data is expected to be accessed frequently, either as a public dataset or in an image analysis workflow, the upfront conversion time to Zarr will lead to overall time savings over frequent accesses. In situations where object storage is mandated as in large scale public repositories, we would encourage the use of OME’s next generation file formats (OME-NGFF) today. Alternatively, users who will need to frequently transmit their images may choose to store their data in a large single file like HDF5. OME-TIFF remains a safe option for those who rely on proprietary software for image viewing and analysis, especially in digital pathology and other whole slide image applications, as many have been extended to both read and write this open standard. Each choice comes with benefits and costs, and individual scientists, institutions, global collaborations and public data resources need the flexibility to decide which approach is appropriate. Regardless, we foresee a fundamental change in work patterns where data generated in advanced bioimaging applications is converted into a format that is optimised for downstream processing, analysis, visualization and sharing, and all subsequent data access occurs via open data formats, without the need for repeated, on-the-fly translation. When conversion of primary imaging data is an option, we would encourage the community to choose from the most appropriate of the formats supported by OME.

Ultimately, we hope to see digital imaging systems producing open, transparent, in others words FAIR, data without the need for further conversion. Until that time, we are committed to providing the data conversion needs of the community. Following the same pattern we have used with *bioformats2raw* and *raw2ometiff*, we propose to meet this challenge via a set of migration tools allowing efficient data transformations between all data formats contained in this suite of interoperable formats. Additionally, as the specification evolves based on community feedback, the same migration tools will allow upgrading the scientific data generated by the bioimaging community to prevent the need for long-term maintenance of older data. Upcoming specifications include geometric descriptions of regions of interest, meshes, transformations for correlative microscopy, and others.

To provide the best chance of wide adoption and engagement, we are developing the formats in the open, with frequent public announcements of progress and releases of reference software and examples (https://forum.image.sc/tag/ome-ngff) and regular community meetings where we present work, source feedback, and encourage community members, including vendors, to participate in the specification and implementation. The community process is being developed and we welcome contributions from all interested parties on https://github.com/ome/ngff.

## Acknowledgements

Work on the *bioformats2raw* and *raw2ometiff* converters was funded by awards from InnovateUK to Glencoe Software Ltd (PathLAKE, Ref: 104689 and iCAIRD Ref: 104690). Work on next generation file formats by OME was funded by grant number 2019-207272 from the Chan Zuckerberg Initiative DAF, an advised fund of Silicon Valley Community Foundation, the Wellcome Trust (Ref: 212962/Z/18/Z) and BBSRC (Ref: BB/R015384/1).

The authors would like to thank the originators of the Zarr and N5 formats, Alistair Miles and Stephan Saalfeld, and the vibrant communities they have built for working together to unify their formats.

The development of OME-NGFF has been and will continue to be a community endeavor. Everyone who has participated in the format specification and/or an implementation is invited to request software authorship (http://credit.niso.org/contributor-roles/software/) by contacting the corresponding author.

